# Pheromone binding protein shapes olfactory temporal resolution

**DOI:** 10.1101/2021.01.11.426239

**Authors:** Yusuke Shiota, Takeshi Sakurai, Noriyasu Ando, Stephan Shuichi Haupt, Hidefumi Mitsuno, Takaaki Daimon, Ryohei Kanzaki

**Affiliations:** Research Center for Advanced Science and Technology, The University of Tokyo, 4-6-1 Komaba, Meguro-ku, Tokyo 153-8904, Japan; Department of Agricultural Innovation for Sustainability, Faculty of Agriculture, Tokyo University of Agriculture, 1737 Funako, Kanagawa, Atsugi 243-0034, Japan; Department of Applied Biosciences, Graduate School of Agriculture, Kyoto University, Kitashirakawa Oiwakecho, Sakyo-ku, Kyoto 606-8502, Japan

## Abstract

Male moths are capable of orienting towards conspecific females using sex pheromones. Since pheromones are distributed as discontinuous plumes owing to air turbulence, tracking intermittent stimuli with high temporal resolution is suggested to be important for efficient localisation. Here, using a pheromone binding protein (BmPBP1) knockout silkmoth, we revealed that the loss of functional pheromone binding protein altered antennal response kinetics resulting in reduced temporal resolution to intermittent pheromone stimuli on the antennae. Behavioural analysis revealed that *BmPBP1*-knockout males exhibited significantly less straight walking, which occurs when detecting pheromone stimuli, especially to high frequency stimuli. Accordingly, *BmPBP1*-knockout males took a significantly longer time to locate pheromone sources and females than did wild-type males. Together, BmPBP1 plays a critical role in determining temporal antennal response kinetics and that an appropriate range of temporal sensory and behavioural resolutions is essential for tracking pheromone plumes for efficient pheromone source localisation in the silkmoth.

## Introduction

Many animals rely on olfaction to locate potential mating partners or food sources for survival in a natural environment. Odour source localisation is a difficult task for animals because odours are distributed discontinuously owing to turbulent air flow in the natural environment; the stochastic nature of odour distribution further increases the unpredictability of odour arrival^1-3^. Mechanisms underlying how animals can accurately locate odour sources in such a complex environment have long been a core topic of study in the field of olfactory research.

The sex pheromone communication system in moths is one of the best examples of sophisticated odour source localisation in animals, whereby male moths detect and orient toward conspecific females using intermittent sex pheromone information emitted by females. Owing to the simple relationship between sex pheromone chemicals and the localisation behaviour, pheromone source localisation behaviours can be relatively easily induced in the laboratory. As such, this behaviour has been studied as an essential model system to better understand the general mechanisms underlying odour source localisation in animals.

Odour source localisation behaviours of male moths consist of two basic behavioural components: an upwind surge and crosswind casting^4^. The surge is a straight upwind flight triggered shortly after exposure to the pheromone filament, whereas crosswind casting is a self-steering counterturning that occurs when a male moth loses contact with the pheromone filaments in the plume. In the silkmoth *Bombyx mori*, when exposed to the pheromone, male moths exhibit straight walking at a small turning angle as a reflex behaviour (surge), followed by left and right zigzag turns (casting), with a gradually increasing turn angle when they encounter an air pocket of a pheromone plume^5-7^. This behavioural sequence is reset in response to every new exposure to a pheromone stimulus. Thus, search-related walking paths are highly dependent on the temporal structure of the pheromone stimuli^5,8,9^. When the stimulation frequency is low, males walk in complex zigzag paths, whereas a higher frequency results in successive surge behaviours and straighter paths. Therefore, male antennae should possess a sufficient temporal resolution to track intermittent pheromone stimulation. The surge-casting (phasic-tonic) behavioural model for odour source localisation is thought to be a common strategy among moth species underlying their localisation behaviour^4,10^.

The importance of intermittent odour information has also been highlighted in experiments using homogeneous pheromone stimulation that rarely contains fluctuating pheromone filaments in the plume (referred to as a homogeneous pheromone cloud). In homogeneous pheromone clouds, moths cannot sustain an upwind surge and exhibit a casting flight, leading to the inability to localise the pheromone source^11,12^. Later, to investigate the physiological responses to intermittent stimuli, the characteristics of sensory output to various odour frequencies, including a homogeneous pheromone cloud, were examined in the moths *Grapholita molesta* and *Heliothis virescens*^10,13^. In response to intermittent stimuli, the electroantennogram (EAG) responses, which monitor the total activity of all olfactory receptor neurons (ORNs) in the antenna, of moths reflected intermittent exposure to pheromone filaments in the plume. However, in a homogeneous pheromone cloud, the responses did not show dynamic changes. These reports conclude that moths naturally receive intermittent pheromone information on their antennae, and this modulates surge-casting localisation behaviour. Therefore, resolving the temporal properties of odour information continuously is thought to be a very important aspect of odour source localisation^14,15^. However, the temporal sensory and behavioural resolution required for successful odour source localisation and the underlying neural and molecular mechanisms remain to be elucidated.

Pheromone binding proteins (PBPs) are small soluble proteins highly enriched in the sensillum lymph space^16^. These proteins are reported to contribute to the sensitivity and/or selectivity of antennal responses^17-21^. Also, PBPs are thought to be involved in the early inactivation of pheromones, following pheromone detection; thus, PBPs are a candidate molecular factor for controlling temporal antennal responses^22-25^. However, this hypothesis has not been experimentally tested yet. In the silkmoth *B. mori*, one of three PBP genes named *BmPBP1*, which is expressed in accessory cells surrounding ORNs and secreted into the sensillum lymph of pheromone sensitive sensilla, is reported to play an important role in pheromone detection^26-28^. In a previous study, we established a *BmPBP1*-knockout silkmoth line and showed that the loss of BmPBP1 affected the sensitivity of antennal responses to pheromone components^21^. In this study, using the *BmPBP1*-knockout silkmoth, we revealed that the temporal antennal resolution of *BmPBP1*-knockout male antennae to intermittent pulse trains was significantly reduced. In addition, *BmPBP1*-knockout male moths took significantly longer to locate pheromone sources owing to lowered temporal behavioural resolution, resulting in inefficient source localisation. Based on our results, we will discuss the effects of BmPBP1 loss and the sensory characteristics that govern the efficiency of odour source localisation behaviour.

## Results

### Time constant of antennal responses was altered in *BmPBP1*-knockout male moths

In a previous study, we reported that antennal responses of *BmPBP1*-knockout silkmoths to pheromone components were significantly lower than those of wild-type moths, but antennae retained a clear responsiveness to pheromone components (bombykol and bombykal)^21^. Using *BmPBP1*-knockout males, we first tested whether PBPs were also involved in the temporal properties of antennal responses by investigating the response and recovery time constants of EAG responses of *BmPBP1*-knockout male moth antennae to single pulse stimulation with sex pheromone components (Fig. 1a). Comparison of these parameters with the same bombykol dosage (1000 and 10000 ng) revealed that there were significant differences in the kinetic parameters consisting of latency and recovery time constant at 10000 ng between *BmPBP1*-knockouts and wild-types (Fig. 1b top, 1c). Because the peak EAG amplitudes of *BmPBP1*-knockout male antennae were significantly lower than those of wild-type antennae (Fig. 1b top)^21^, we hypothesised that differences in peak amplitudes could affect the temporal parameters of EAG responses. When we normalised the temporal parameters using peak amplitudes, we found that recovery time constants showed a positive correlation with peak amplitudes, and, as a result, normalised recovery time constants of *BmPBP1*-knockout male antennae were significantly longer than those of wild-type male antennae (p=0.0014 for bombykal; p=0.0017 for bombykol) (Fig. 1b bottom, 1d). However, the other two parameters did not show any correlation with peak amplitude, and no significant differences were found in the response time constant or latency of antennal responses of *BmPBP1*-knockout and wild-type males (Fig. 1b bottom, 1d). These differences were observed with both bombykol and bombykal (Fig. 1d), suggesting that the loss of BmPBP1 delayed the recovery time constant of EAG responses with both bombykol and bombykal.

**Figure 1.**
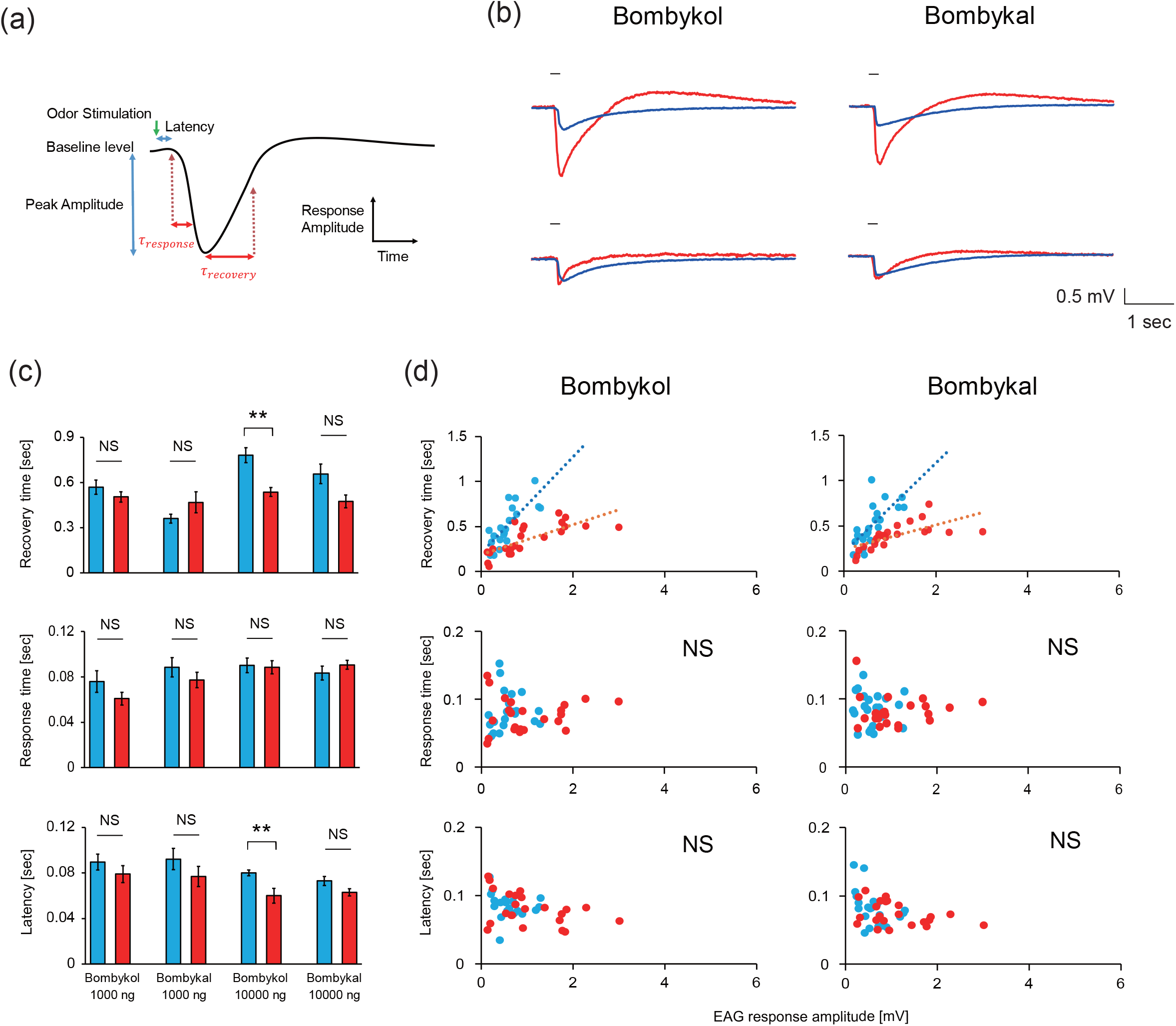
Antennal response kinetics in *BmPBP1*-knockout male moths. **(a)** Parameters considered in the antennal response kinetics analysis of EAG recordings. **(b)** Representative EAG responses of *BmPBP1*-knockout (blue) and wild-type (red) male antennae to 10000 ng bombykol stimulation (top), and responses to 10000 ng bombykol stimulation in *BmPBP1*-knockout (blue) and 100 ng bombykol stimulation in wild-type (red) male antennae (bottom). The stimulus was applied for 200 ms, as indicated by the solid line on the trace. **(c)** Kinetic analysis of EAG responses in *BmPBP1*-knockout (blue; n=11) and wild-type (red; n=5) male antennae. Error bars represent ± SEM. The asterisks indicate significant differences between the groups (\*\**p* < 0.01), as determined using Student’s *t*-test for comparing pairs of data. **(d)** Kinetic analysis of EAG responses of *BmPBP1*-knockout (blue; n=11) and wild-type (red; n=5) antennae normalised by peak EAG amplitude. Broken lines indicate linear regression curves. An ANCOVA test was used to detect significant differences between wild-type and *BmPBP1*-knockout datasets for recovery time, while significant differences between the groups were not detected in the other two parameters (\*\**p* < 0.01). NS indicates not significant.

### Kinetics of temporal antennal responses to pheromone pulse trains in *BmPBP1*-knockout moths

Because tracking intermittent pheromone pulse trains is important to locate pheromone sources in male moths, we next assessed the effects of loss of BmPBP1 on temporal antennal responses to pheromone pulse trains. Particularly, we focused on responses to bombykol because only bombykol triggers the pheromone source localisation behaviour of male silkmoths. We tested three different frequencies of pulse trains, consisting of 0.17 Hz as the low stimulus frequency, 0.83 Hz as the representative of the emission rate of pheromone from females^29^ and 2 Hz as the frequency that can induce a consecutive surge in male silkmoths^5^ (Fig. 2a). Consistent with single pulse experiments, the recovery time constant of EAG responses became longer in *BmPBP1*-knockout than in wild-type male antennae (Fig. 2a). The time required to return to baseline levels after the first bombykol stimulation (termination time) was significantly longer in *BmPBP1*-knockout than in wild-type male moths with all tested frequencies (Fig. 2b, 2c). More importantly, the termination times got longer as the frequency increased and EAG responses of *BmPBP1*-knockout never returned to baseline levels during high frequency stimuli in several tested antennae (1 out of 6 tested antennae at 0.83 Hz, 4 out of 12 tested antennae at 2 Hz). However, all EAG responses of wild-type male antennae quickly recovered to baseline levels (n=7 at 0.83 Hz, n=6 at 2 Hz) (Fig. 2c).

**Figure 2.**
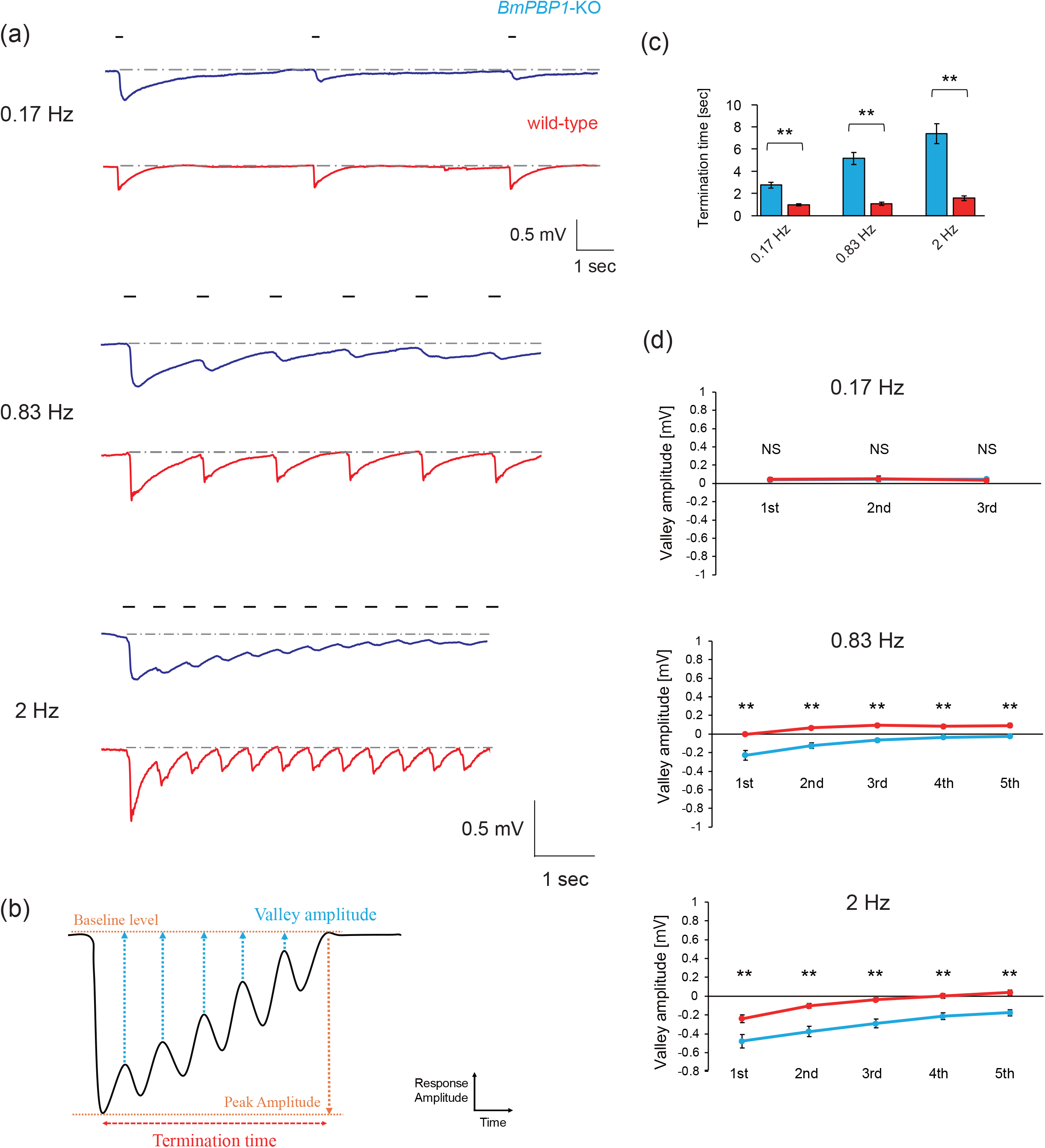
Analysis of antennal response kinetics to pulsed pheromone stimuli in *BmPBP1*-knockout male moths. **(a)** Representative EAG responses to pulsed pheromone trains in *BmPBP1*-knockout (blue) and wild-type (red) male antennae at 0.17 Hz, 0.83 Hz and 2 Hz. The stimulus was applied for 200 ms, as indicated by the solid line on the trace. The broken line indicates baseline level of each EAG trace. **(b)** Parameters considered in the antennal response kinetics analysis of EAG recording to pulsed pheromone stimuli. **(c)** Termination time of EAG responses to pulsed pheromone trains in *BmPBP1*-knockout (blue) and wild-type (red) antennae at 0.17 Hz (KO; n=9, WT; n=9), 0.83 Hz (KO; n=6, WT; n=7) and 2 Hz (KO; n=12, WT; n=6). Error bars represent ± SEM. The asterisks indicate significant differences between the groups (\*\**p* < 0.01), as determined using Student’s *t*-test for comparing pairs of data. **(d)** Valley amplitude of male antennae to pheromone stimuli in *BmPBP1*-knockout (blue) and wild-type (red) at 0.17 Hz (KO; n=9, WT; n=9), 0.83 Hz (KO; n=7, WT; n=7) and 2 Hz (KO; n=16, WT; n=7). The asterisks indicate significant differences between the groups (\*\**p* < 0.01), as determined using Student’s *t*-test for comparing pairs of data. NS indicates not significant.

To evaluate response recovery after intermittent stimuli, we calculated the valley amplitudes, which were defined as the EAG amplitude where each local minimum amplitude was subtracted from the baseline level amplitude (Fig. 2b, 2d, see also Methods section). The valley amplitudes of *BmPBP1*-knockout males were significantly larger than those of wild-type males when stimuli were delivered at 0.83 and 2 Hz. In comparison, the valley amplitudes of both *BmPBP1*-knockout and wild-type males returned to baseline levels after each stimulus was delivered at 0.17 Hz, and there was no significant difference between the groups (Fig. 2d). These results indicate that delayed recovery time constant in *BmPBP1*-knockout male antennae affects the overall temporal response kinetics of EAG responses to intermittent pheromone stimuli.

### Temporal resolution of a single ORN in *BmPBP1*-knockout male moths was lower compared to that in wild-type male moths

To understand the physiological basis of the observed temporal characteristics in EAG experiments, we next performed single sensillum recordings (SSRs) of pheromone sensitive sensilla trichodea (Fig. 3a). We revealed that spontaneous spike activity was not significantly different between *BmPBP1*-knockout and wild-type antennae, suggesting that BmPBP1 does not affect spontaneous spike levels (Fig. 3b, 3c). To compare temporal kinetics, we used 10000 ng bombykol as stimuli for *BmPBP1*-knockout male antennae and 1000 ng bombykol as stimuli for wild-type male antennae, which evoked similar spike counts in *BmPBP1*-knockout male antennae and wild-type male antennae (Fig. 3b, 3c). The spike responses of *BmPBP1*-knockout male antennae at 0.17 Hz took significantly longer to return to spontaneous spike levels, while those of wild-type antennae quickly terminated after the stimulus (Fig. 3d). When antennae were stimulated at 2 Hz, this tendency became clearer, and the spike responses of all tested ORNs in *BmPBP1*-knockout antennae did not return to spontaneous spike levels between subsequent stimuli, while wild-type ORN spike responses quickly returned to spontaneous levels (Fig. 3e, 3f). These results demonstrate that the loss of BmPBP1 delayed response termination of ORNs resulting in a lower antennal temporal resolution in *BmPBP1*-knockout males.

**Figure 3.**
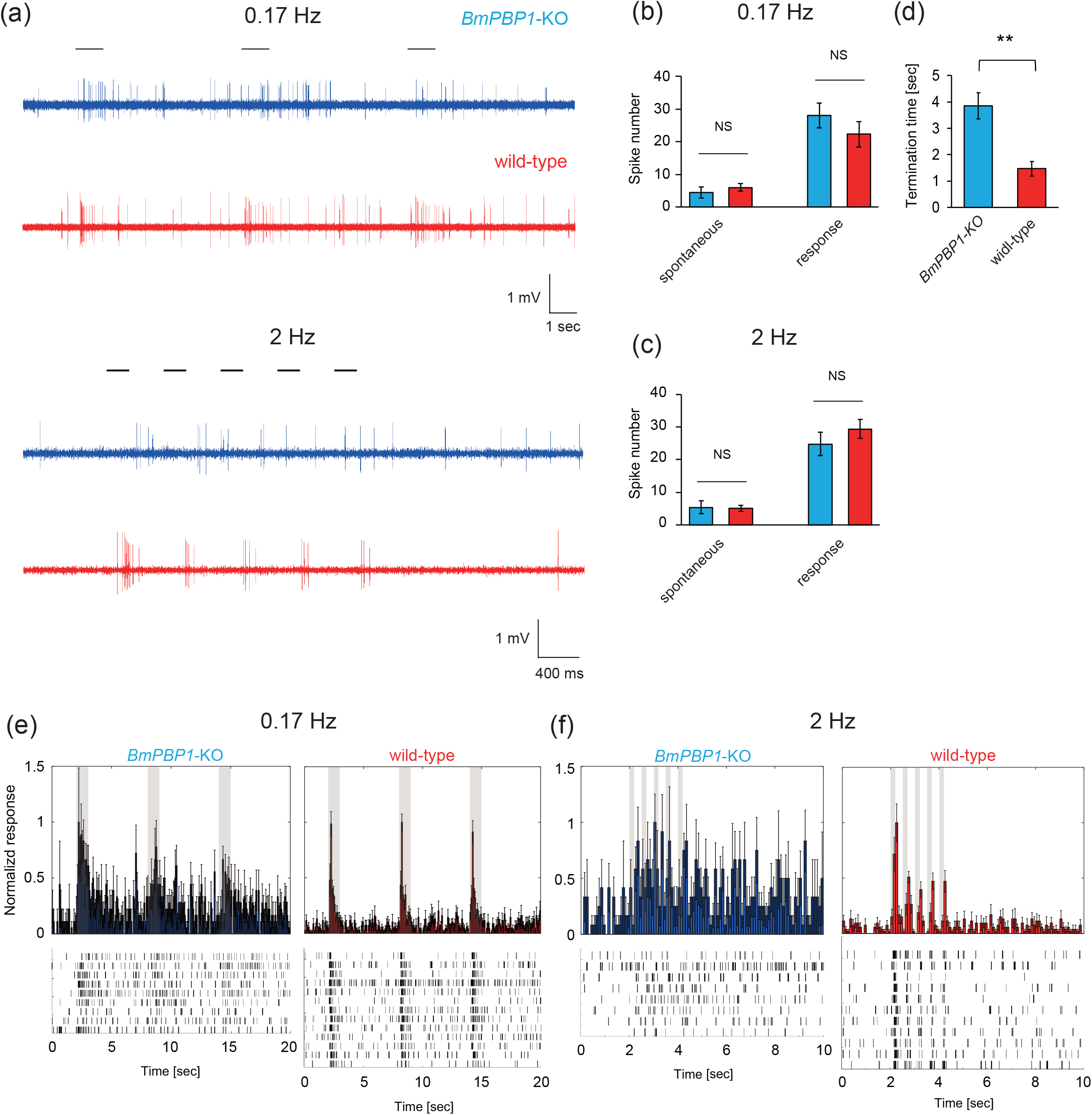
Analysis of antennal temporal resolution to pheromone pulse trains in *BmPBP1*-knockout male moths. **(a)** Representative spike responses of long sensilla trichodea in *BmPBP1*-knockout (blue) and wild-type (red) male antennae to pheromone pulse stimuli at 0.17 Hz and 2 Hz. Solid lines on the trace represent the timing of intermittent bombykol stimuli. **(b)** Comparison of spontaneous and elicited spike numbers. Spontaneous spikes were calculated with a time bin of 2 seconds before the first stimulus was presented to *BmPBP1*-knockout (blue; n=9 at 0.17 Hz, n=8 at 2 Hz) and wild-type (red; n=13 at 0.17 Hz, n=11 at 2 Hz) male antennae; spike numbers were calculated with a time bin of 5 seconds from the stimulus at 0.17 Hz and **(c)** with a time bin of 3.5 seconds after the stimulus at 2 Hz. Error bars represent ± SEM. The asterisks indicate significant differences between the groups (\*\**p* < 0.01), as determined using Student’s *t*-test for comparing pairs of data. **(d)** Comparison of the termination time of spikes between *BmPBP1*-knockout and wild-type at 0.17 Hz. Error bars represent ± SEM. The asterisks indicate significant differences between the groups (\*\**p* < 0.01), as determined using Student’s *t*-test for comparing pairs of data. **(e)** Spike rate in time bins of 100 ms in *BmPBP1*-knockout (blue; n=9) and wild-type (red; n=13) male antennae at 0.17 Hz and **(f)** those of 100 ms in *BmPBP1*-knockout (blue; n=8) and wild-type (red; n=11) male antennae at 2 Hz. Error bars represent ± SEM. The asterisks indicate significant differences between the groups (\**p* < 0.05, \*\**p* < 0.01), as determined using Student’s *t*-test for comparing pairs of data. NS indicates not significant.

### Temporal behavioural resolution of *BmPBP1*-knockout male moths was lower compared to that of wild-type male moths

To investigate the effects of delayed termination kinetics on pheromone source localisation behaviours (Fig. 4a), we first analysed pheromone source localisation behaviours to bombykol using tethered male moths. Upon exposure to a single pulse of bombykol, each *BmPBP1*-knockout male moth exhibited full pheromone-induced localisation behaviour, consisting of a surge behaviour followed by zigzag turns (Fig. S1, Fig. 4b), indicating that the loss of BmPBP1 did not abolish innate behavioural sequences, despite the surge duration of *BmPBP1*-knockout male moths being significantly longer than that of wild-type male moths (Fig. 4b).

**Figure 4.**
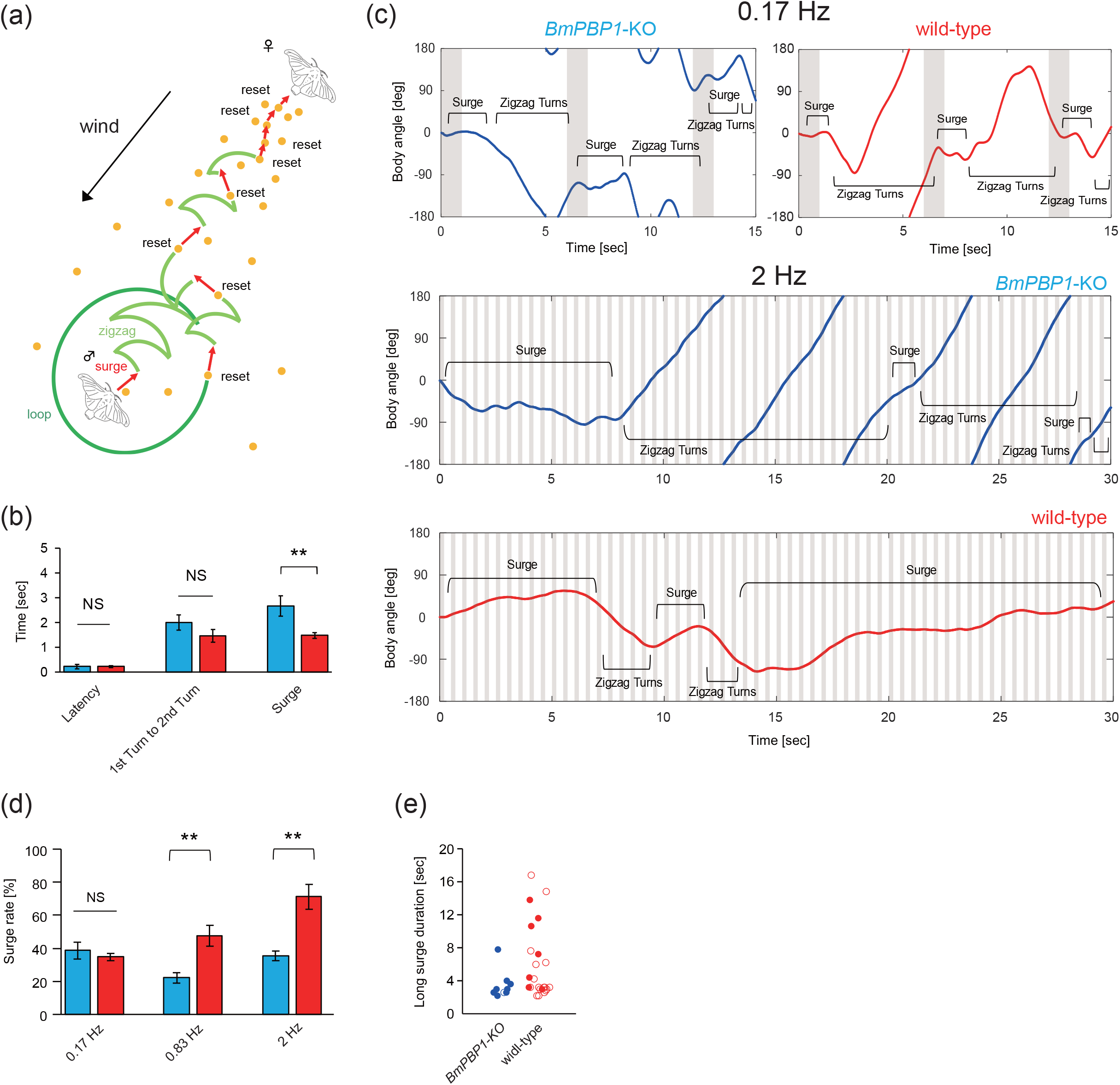
Behavioural responses of tethered *BmPBP1*-knockout moths to intermittent pheromone pulses. **(a)** Schematic diagram of the pheromone-triggered programmed behaviour and pheromone source localisation strategy of a male silkmoth, *B. mori*. **(b)** Analysis of pheromone-triggered programmed behaviour in *BmPBP1*-knockout (blue; n=9) and wild-type (red; n=16) male moths to single pulsed stimulation. Bombykol pulse stimuli were applied for 1 second. Error bars represent ± SEM. The asterisks indicate significant differences between the groups (\*\**p* < 0.01), as determined using Student’s *t*-test for comparing pairs of data. **(c)** Representative trace of the body angle of tethered *BmPBP1*-knockout (blue) and wild-type male (red) moths to pulsed bombykol stimuli at 0.17 Hz and 2 Hz. **(d)** Surge rate in *BmPBP1*-knockout (blue) and wild-type (red) male moths to bombykol pulse trains at 0.17 Hz (KO; n=9, WT; n=16), 0.83 Hz (KO; n=10, WT; n=7) and 2 Hz (KO; n=9, WT; n=9). Error bars represent ± SEM. The asterisks indicate significant differences between the groups (\*\**p* < 0.01), as determined using Student’s *t*-test for comparing pairs of data. **(e)** Scatter plot of each surge based on surge duration in *BmPBP1*-knockout (blue; n=9) and wild-type (red; n=9) male moths. Circles indicate the distribution of individual data. Solid circles indicate the first surge that was induced by the first bombykol stimuli. NS indicates not significant.

If *BmPBP1*-knockout males perceived intermittent stimuli as continuous signals owing to delayed response kinetics, we hypothesised that behavioural patterns to pulse trains should be changed, especially in terms of resetting behaviour sequences. In response to 0.17 Hz intermittent bombykol stimulation, to which *BmPBP1*-knockout antennae return to baseline levels, both *BmPBP1*-knockout and wild-type male moths showed a surge in exposure to each bombykol pulse (Fig. 4c, d), indicating that each pulse induced the resetting of behaviour sequences. However, following 0.83 Hz and 2 Hz bombykol pheromone pulse trains, *BmPBP1*-knockout male moths displayed lower surge rates—the ratio of surge duration against total locomotion duration (see Methods)—and higher zigzag turn rates than wild-type male moths (Fig. 4c, d). Further, *BmPBP1*-knockout male moths exhibited fewer longer surges (defined as a surge lasting more than 2 seconds) than wild-type male moths (Fig. 4e), indicating that *BmPBP1*-knockout males failed to exhibit successive surge behaviours in response to intermittent pheromone stimuli at 2 Hz. Importantly, differences in surge rates were well correlated with EAG and SSR results; males restarted surge behaviours to each pheromone stimulus under stimulus conditions that allowed antennal responses to return to baseline levels, presumably owing to failure to clearly detect the onset of the subsequent stimulus. Thus, our results indicate that lower temporal antennal resolution reduces temporal behavioural resolution in *BmPBP1*-knockout males.

### Wind tunnel experiments revealed behavioural deficits in efficient pheromone source localisation in *BmPBP1*-knockout male moths

To assess whether changes in the behavioural patterns of *BmPBP1*-knockout males affect odour source localisation, we investigated pheromone source localisation behaviours to 2 Hz pheromone pulse trains in a wind tunnel under free walking conditions. We analysed two criteria for efficient pheromone source localisation: success rate and localisation time. To compensate for differences in EAG sensitivity (Fig. 1) and to test the effects of pheromone dose on pheromone source localisation, the behaviours of *BmPBP1*-knockout and wild-type males were tested with 100–10000 ng and 1–100 ng bombykol, respectively. Both *BmPBP1*-knockout and wild-type males could localise the pheromone source for all tested doses of bombykol with a similar success rate (Fig. 5a). However, *BmPBP1*-knockout males took significantly longer to localise the pheromone source than wild-type males (Fig. 5b). Therefore, though *BmPBP1*-knockout males still possess the ability to localise the pheromone sources, the efficiency of their localisation behaviour in terms of localisation time was significantly reduced in *BmPBP1*-knockout males.

**Figure 5.**
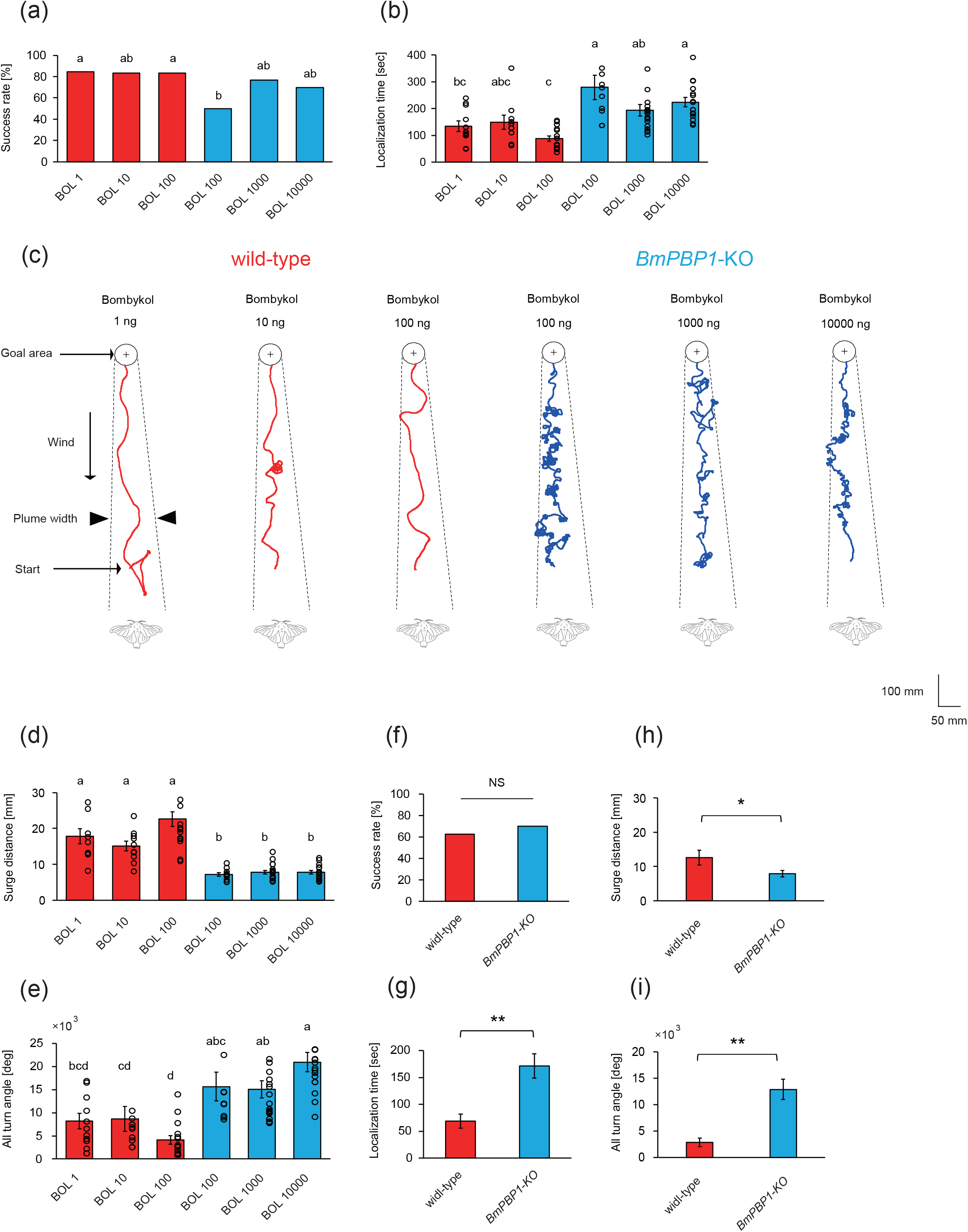
Performance of pheromone source localisation in *BmPBP1*-knockout male moths. **(a)** Success rate of pheromone source localisation of *BmPBP1*-knockout (blue) and wild-type (red) male moths at 2 Hz (WT; n=13 at 1 ng; n=12 at 10 ng; n=18 at 100 ng, KO; n=20 at 100 ng; n=26 at 1000 ng; n=23 at 10000 ng). The different letters indicate significant differences according to Fisher’s exact probability test. **(b)** Pheromone source localisation time of *BmPBP1*-knockout (blue) and wild-type (red) male moths at 2 Hz (KO; n=10 at 100 ng; n=20 at 1000 ng; n=17 at 10000 ng, WT; n=11 at 1 ng; n=10 at 10 ng; n=15 at 100 ng). Error bars represent ± SEM. The different letters indicate significant differences according to the Steel–Dwass test. Black solid circles indicate the distribution of individual data. **(c)** Representative trajectory of *BmPBP1*-knockout (blue) and wild-type male (red) moths to tested bombykol dosage (WT: 1, 10, 100 ng, KO: 100, 1000, 10000 ng) during pheromone source localisation. Bombykol pulse stimuli were applied for 200 ms at 2 Hz. Solid lines indicate the trajectory of the pheromone source localisation behaviour. Broken lines indicate the estimated boundaries of the pheromone plume. **(d)** Surge distance of *BmPBP1*-knockout (blue) and wild-type (red) male moth to bombykol pulse trains at 2 Hz (WT; n=11 at 1 ng; n=10 at 10 ng; n=15 at 100 ng, KO; n=10 at 100 ng; n=20 at 1000 ng; n=17 at 10000 ng). **(e)** All turn angles of *BmPBP1*-knockout (blue) and wild-type (red) male moths to bombykol pulse trains at 2 Hz (WT; n=11 at 1 ng; n=10 at 10 ng; n=15 at 100 ng, KO; n=10 at 100 ng; n=20 at 1000 ng; n=17 at 10000 ng). **(f-i)** Performance of pheromone source localisation to a female moth. Success rate (WT; n=10, KO; n=8), localisation time (WT; n=7, KO; n=5), surge distance (WT; n=7, KO; n=5), and all turn angles (WT; n=7, KO; n=5) to a female moth. The asterisks indicate significant differences between the groups (\*\**p* < 0.01), as determined using Student’s *t*-test for comparing pairs of data. NS indicates not significant.

The walking traces of *BmPBP1*-knockout males had distinct patterns with more turning and less successive surge behaviour compared with wild-type males (Fig 5c). Indeed, a detailed analysis of the traces revealed that *BmPBP1*-knockout males exhibited significantly shorter surge distances and a larger summed turning angle during localisation than wild-type males (Fig. 5d, e). Exposure to relatively high bombykol concentrations in *BmPBP1*-knockout male moths (KO: 1000–10000 ng) and low bombykol concentrations in wild-type male moths (WT: 1–10 ng) did not alter the characteristics of localisation behaviours (Fig. 5a-e). Lastly, we examined localisation behaviour to female silkmoths and found that behavioural parameters including success rate (Fig. 5f), localization time (Fig. 5g), surge distance (Fig. 5h) and all turn angles (Fig. 5i) showed the same characteristics with those observed to bombykol. Taken together, these results indicate that behavioural deficits caused by the low temporal antennal resolutions of male moths are responsible for the inefficiency of odour source localisation in *BmPBP1*-knockout males.

## Discussion

In this study, we demonstrated that the temporal sensory characteristics of antennae are crucial for efficient pheromone source localisation and that BmPBP1 is involved in shaping the temporal sensory characteristics of antennae to pheromone stimulation in *B. mori*. Since the discovery of PBPs in 1981 in *Antheraea polyphemus*^16^, PBPs have been suggested to play important roles in various aspects of pheromone detection in moths. Although previous studies have provided evidence that PBPs can enhance the sensitivity of antennal response to pheromones^20,21^, no study has provided *in vivo* experimental evidence for the contribution of PBPs for improving temporal sensory resolution, one of the most important aspects in odour source localisation under natural conditions. Our physiological and behavioural analyses using *BmPBP1*-knockout silkmoths present the first evidence supporting the functional role of BmPBP1 in determining the temporal kinetics of pheromone responses.

Odour source localisation utilises the natural intermittency of odour stimulation to promote a surge when a stimulus pulse is encountered. This behavioural strategy for odour source localisation is, at present, widely accepted in moth species, based on the correspondence between physiological antennal responses and surge behaviours during localisation^10,13^. Therefore, moths require mechanisms that allow for the faithful representation of intermittent odour information at the level of the antennal receptors, as well as a strategy to reflect sensory information in behaviours with sufficient temporal resolution. In this study, we showed that *BmPBP1*-knockout males had a reduced temporal antennal resolution compared with wild-type, resulting in the inefficient localisation to odour sources. Even for the localisation to female silkmoths, similar behavioural deficits, such as a shorter surge distance, longer turning behaviour and longer localisation time, were observed in *BmPBP1*-knockout males. Therefore, these results indicate that moths possess an appropriate range of temporal antennal and behavioural resolutions to efficiently localise odour sources in a dynamically changing environment. Though sensitivity of *BmPBP1*-knockout antennae is lower than that of wild-type antennae, the behavioural deficits are thought to be caused by the alternation of temporal kinetics, but not by the lowered sensitivity, as the behavioural deficits were observed in response to pheromone stimuli that induced similar EAG peak amplitudes in both *BmPBP1*-knockout and wild-type antennae.

Our EAG results demonstrated that the recovery time of EAG responses of *BmPBP1*-knockout male antennae was significantly longer than those of wild-type male antennae, which resulted in the lower temporal resolution of EAG responses to pulsed pheromone stimulation. Although EAG is thought to monitor the total activity of all ORNs in the antenna, the actual relationship between EAG responses and the spike activity of individual ORNs is not clear. Thus, it is important to note that consistent temporal characteristics were also observed at the level of single ORN spike responses, showing that EAG response kinetics in *BmPBP1*-knockout antennae reflects changes in temporal spike characteristic of ORNs.

Similar temporal alterations in antennal and behavioural responses to odour pulse trains have been previously reported in the oriental fruit moth *Grapholita molesta*^30,31^. Under cool temperature conditions, antennal responses of *G. molesta* males to pheromones were reduced, and they were unable to keep up with high frequency pulsed pheromone stimuli due to the inability to generate well-contrasted on-off spike representations to each stimulus. This characteristic property of the spike response of ORNs is called “attenuation”^30^. Under similar cooling conditions, male moths failed to initiate successive surge behaviours and could not locate an odour source^31^. Although the underlying mechanism that affects the antennal response characteristics and behaviours may be different, these studies also indicate the importance of the temporal characteristics of antennal responses for efficient pheromone source localisation. In our study, we revealed that BmPBP1 is one factor related to temporal antennal resolution and that temporal sensory representations corresponding to odour dynamics play a key role in the quick behavioural reset strategy for efficient localisation.

We found that the stimuli frequencies to which EAG responses could recover from peak amplitude to baseline levels corresponded to those where behavioural sequences of male moths were reset. Therefore, after peak neuronal activity, returning to spontaneous baseline activity can be important for tracking odour filaments by continuously resetting behavioural sequences, rather than only peak amplitude detection. Previous studies have reported the temporal antennal resolution of several moth species based on the peak frequency analyses of EAG recordings^32,33^. According to these reports, antennae of male *B. mori* have a potential temporal resolution to intermittent odour pulses of up to 25 Hz. However, since EAG responses to high frequency pheromone stimulation in those reports do not recover to baseline levels after peak response, there is a possibility that not every odour stimulus is resolved to induce resetting behavioural sequences and initiation of surge behaviour. Thus, the potential temporal resolution of antennae may differ from the antennal temporal resolution for a behavioural reset. In other words, sensory responses may turn into ‘fused’ information for male moth behaviour when tracking an odour plume at a frequency where neural activity does not return to spontaneous baseline activity before a response to a subsequent odour stimulus. In the future, investigation of the behavioural patterns of wild-type animals to high frequency stimuli that induce antennal responses without returning to baseline levels can shed light on antennal response parameters crucial for the reset of behaviour sequences.

Previous kinetics model analyses proposed that rapid inactivation of pheromone molecules after they activate pheromone receptors plays a critical role in controlling the kinetics of pheromone responses and that PBPs were one candidate molecular component of pheromone inactivation^23-25^. Our results showing that the loss of BmPBP1 delayed recovery time constants strongly support the idea that BmPBP1 is involved in the rapid inactivation of pheromone molecules. PBPs have shown to be involved in sensitive detection of pheromones by efficiently solubilising pheromone molecules into the sensillum lymph and protecting them from enzymatic degradation during transportation to the vicinity dendritic membrane of ORNs, where pheromone receptors resides^21,22^. Therefore, PBPs play multiple functional roles both before and after pheromone detection by pheromone receptors. Further electrophysiological, biochemical and molecular analyses of *BmPBP1*-knockout moths will help reveal the detailed mode of action underlying how PBPs can perform these different functions in the process of pheromone detection.

In this study, we reported that BmPBP1 contributes to the temporal control of antennal responses, and its absence lowered the antennal temporal resolution, critically affecting efficient odour source localisation. Our results could provide a new methodology for controlling pest insects by modulating odour source localisation behaviour. Furthermore, strategies to represent spatio-temporal airborne odours in the sensory system by quickly terminating signals after detection, and to reset sequential behavioural patterns in response to sensory input, are thought to be equally effective for the development of artificial odour source searching algorithms.

## Methods

### Animals and chemicals

We used *BmPBP1*-knockout moths, which were generated using transcription activator-like effector nuclease (TALEN)-mediated gene targeting, as described in a previous study^21^. Larvae were reared on an artificial diet (Nihon Nosan Kogyo, Yokohama, Japan) at 25°C under a 16:8 h (light/dark) photoperiod. Following eclosion, one to seven day old male moths were used for the experiments. Subsequent to all experiments in this study, the genotype of all *BmPBP1*-knockout males was determined by PCR, as previously described^21^. The purity (>99.5%) of synthetic bombykol and bombykal was verified by gas chromatography using previously described conditions^34,35^, and they were kindly provided by Dr. S. Matsuyama from the University of Tsukuba, Japan.

### Electroantennogram (EAG) recordings

An antenna of a male moth was excised at the base, and a few antennomeres at the tip were cut off. The antenna was then mounted on the EAG probe using electrode gel (SPECTRA 360; Parker Laboratories, Fairfield, NJ, USA). A glass cartridge (inner diameter, 5 mm) was prepared for stimulation by inserting a piece of filter paper (1.5 × 1.5 cm), and 5 μL of a pheromone solution in *n*-hexane was administered. A charcoal-purified airstream (1 L/min) was passed through the glass cartridge and directed onto the antenna. The EAG responses were amplified using a custom-made amplifier (Minegishi and Kanzaki, unpublished), band-pass filtered at 0.16 to 300 Hz, and digitised at 1 kHz (USB-6210; National Instruments, Austin, TX, USA). The data were analysed using a custom-written program (MATLAB; Mathworks, Natick, MA, USA). Baseline levels were calculated from 2 sec of data recording before the stimulus. The start time of the response was defined as the time the EAG amplitude exceeded the baseline level by ± 2.5 standard deviations. The response time constant was defined as the time of the response phase, from the start of response to 63.2 % of the absolute EAG response amplitude. The recovery time constant was defined as the time of the settling phase, from the peak amplitude to 63.2% of the absolute EAG response amplitude. The valley amplitude was defined as the EAG response amplitude where each local minimum amplitude was subtracted from the baseline level. The termination time was defined as the time from the peak amplitude to the baseline level by ± 5% standard error of the mean (SEM). The latency was calculated from the start time by subtracting the arrival time of the pheromone on the antenna that was estimated using a method previously described^36^.

### Single sensillum recording (SSR)

Single sensillum recordings were performed in a Faraday cage. Moths were fixed on a custom-made acrylic chamber under an Olympus BX50WI (500×) microscope. The antennae were stabilised with dental wax (GC Corporation, soft plate wax), and the basement segments were fixed with a resin bond (GC Corporation, G-Fix). Spike responses were recorded by a sharpened tungsten wire electrode (diameter 0.5 mm, tip approximately 1 μm) inserted into the bases of pheromone sensitive long sensilla trichodea on the antennae. As a reference electrode, a silver wire was inserted into the compound eye of the moth. Pheromone was delivered by injecting pheromone pulses into a charcoal-purified and moistened continuous airstream (1 L/min), to reduce any mechanical artefacts. Spike responses were band-pass filtered at 300 to 3k Hz and amplified (Nihon Koden, MEZ-8300). All spike responses were digitised at 10 kHz. The spontaneous spike rate was calculated from 2 sec of data recording before the stimulus, with a time bin of 100 ms. We analysed data with a custom MATLAB (Mathworks, MA, Natick, USA) program and Spike2 software (Cambridge Electronic Design Limited, UK).

### Behavioural experiments using tethered moths

Scales on the dorsal thorax of moths were removed, and a copper wire holder was attached to the thorax with glue. Moths were placed on the top of a 70 mm diameter polystyrene foam ball. The ball was supported by a plastic funnel with an electrical fan attached under the ball. The air flow generated by the electric fan kept the ball floating in the air to reduce friction and enable moths to behave during locomotion. Pulsed pheromone stimulation of 200 ms duration at 0.83 Hz and 2 Hz frequency and stimulation of 1 second at 0.17 Hz frequency was used. Wild-type and *BmPBP1*-knockout males were stimulated with 100 ng and 10000 ng bombykol, respectively. Based on a previous study using tethered moths, the surge behaviour was defined as straight walking with a small angle, at an accumulated angular velocity ≤ 8°, when the accumulated walking distance ≥ 1 mm, with a forward velocity ≥ 1 mm/sec, using a time bin of 200 ms. The zigzag turn behaviour was defined as turning behaviour with a large angle, at an accumulated angular velocity ≥ 8°/sec, with an accumulated forward velocity ≤ 10 mm/sec. These parameters were adjusted to remove trackball noise from oscillations caused by stepping of the moth according to a previous study^37^. Pheromone source localisation behaviours of moths were recorded with an optical mouse attached behind the polystyrene ball. Surge rate was defined as the ratio of total surge duration to total locomotion duration containing both surge and zigzag turns. We processed data based on these criteria with a custom MATLAB (Mathworks, MA, Natick, USA) program.

### Wind tunnel experiments

We performed odour source localisation experiments in a wind tunnel (1800×900×300 mm L×W×H) at 25−28°C. Pulsed pheromone stimulation of 200 ms duration at 2 Hz frequency was controlled by an electric valve, and wind velocity was controlled at 0.5−0.7 m/s. For the odour source localisation experiments, both *BmPBP1*-knockout and wild-type male moths were used within 1 to 4 days after eclosion. Synthetic bombykol (1, 10, 100, 1000 and 10000 ng) and wild-type females were used for pheromone stimulation. Moths were placed 50 cm downstream of a pheromone source. The pheromone source localisation behaviour of male moths was captured with a digital video camera at a frame rate of 30 Hz. Surge was defined as straight walking with a small angle, at an angular velocity ≤ 5°/sec, with the accumulated turn angle ≤ 30°/sec, with a forward velocity ≥ 1 mm/sec and a walking distance ≥ 1 mm, using a bin size of 200 ms based on previous studies^38^. Behaviours that did not correspond to these criteria were seen as stopping and switching directions and were not included in our analyses. The other behaviours (at angular velocity ≥ 5°/sec, with accumulated turn angle ≥ 30°/sec) were classified as zigzag turns. We processed image data based on these criteria with customised Java and R scripts (R Development Core Team, 2010).

### Statistical analysis

In our experiments, to assess the statistical differences between wild-type and *BmPBP1*-knockout moths, we used Student’s *t*-test, Fisher’s exact probability test, the Steel–Dwass test and an ANCOVA test. Statistical significance was calculated using Microsoft Excel 2016, a commercial macroprogram (Statcel version 4; Seiun-sya, Japan) and custom VBA and MATLAB (Mathworks, MA, Natick, USA) programs. The error bars shown in the figures represent the SEMs. The asterisks indicate significant differences between the groups (\**p* < 0.05, \*\**p* < 0.01). The different letters indicate significant differences according to the Steel–Dwass test and Fisher’s exact probability test.

## Acknowledgements

We thank Dr. Ryo Minegishi for the development of the EAG amplifier; Mr. Takuya Nakajo and Ms. Junko Tsuchiya and Ms. Akane Kitazono-Itoigawa for technical assistance in rearing silkmoths; and Dr. Shigeru Matsuyama for providing bombykol. This work was supported by a Grant-in-Aid for Young scientists (A) (26712027), Grant-in-Aid for Scientific Research (B), Japan Society for the Promotion of Science (JSPS), Japan, awarded to T.S. and by a Grant-in-Aid for Scientific Research (B) (15H04399), JSPS, Japan, awarded to R.K.

## Author contributions

Y.S., T.S., and R.K. designed the research; Y.S., T.S., N.A., S.S.H., and H.M. performed the research; T.D. contributed new reagents/analytic tools; Y.S. and T.S. analysed the data; and Y.S. and T.S. wrote the paper.

