## Supplemental Figure 1 for "Pheromone binding protein shapes olfactory temporal resolution"

Takeshi Sakurai, Ph.D.

Department of Agricultural Innovation for Sustainability, Faculty of Agriculture, Tokyo University of Agriculture, 1737 Funako, Kanagawa, Atsugi 243-0034, Japan

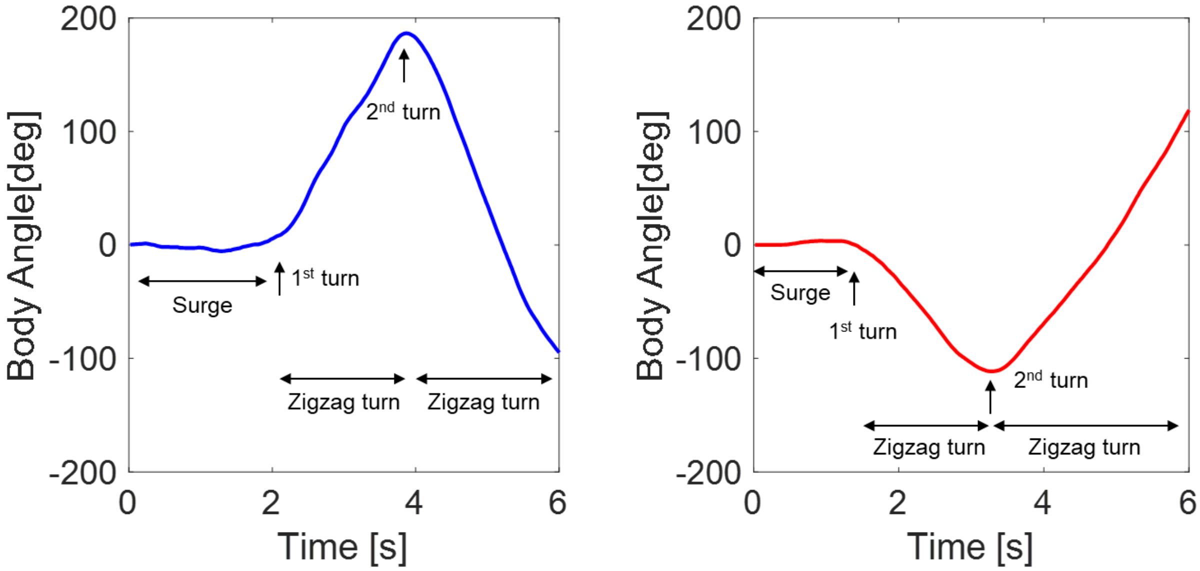


Fig. S1
Representative trace of the body angle of tethered *BmPBP1*-knockout (blue) and wild-type male (red) moths to single bombykol stimulus.
